# Acoustic trapping and navigation of microrobots in the mouse brain vasculature

**DOI:** 10.1101/2023.01.28.522839

**Authors:** Alexia Del Campo Fonseca, Chaim Glück, Jeanne Droux, Yann Ferry, Carole Frei, Susanne Wegener, Bruno Weber, Mohamad El Amki, Daniel Ahmed

**Author notes:** **Correspondence** and requests for materials should be addressed to: Daniel Ahmed, Akustische Robotik Biowiss.u.Med. RSA G 324, Säumerstrasse 4, 8803 Rüschlikon, Switzerland, **E-mail**, Mohamad El Amki, Dept. of Neurology, University Hospital Zurich, Frauenklinikstrasse 26, Zürich, 8091, Switzerland, **E-mail**. **shared last authors** (Mohamad El Amki, Daniel Ahmed). **Author Contributions**: D.A. conceived the project. D.A. and M.E.A conceived and supervised the project. A.C.F performed all the experiments and performed data analysis with contributions from Y.F and C.F and with feedback from D.A and M.E.A. A.C.F, C.G and J.D performed all the experiments *in vivo*. All authors contributed to the experimental design, scientific presentation, and discussion. A.C.F, C.G, M.E.A and D.A. wrote the manuscript. **Competing Interest Statement**: The authors declare no competing financial interests. **Classification**: Physical science – Applied physical science.

## Abstract

Many cerebrovascular and neurodegenerative diseases are currently challenging to treat due to the complex and delicate anatomy of the brain. The use of microrobots can create new opportunities in brain research due to their ability to access hard-to-reach regions and empower various biological applications; however, little is known about the functionality of microrobots in the brain, owing to their limited imaging modalities and intravascular challenges such as high blood flow velocities, osmotic pressures, and cellular responses. Here, we present an acoustic, non-invasive, biocompatible microrobot actuation system, for *in vivo* navigation in the bloodstream, in which microrobots are formed by lipid-shelled microbubbles that aggregate and propel under the force of acoustic irradiation. We investigated their capacities *in vitro* within a microfluidic 3D setup and *in vivo* in a living mouse brain. We show that microrobots can self-assemble and navigate upstream in the brain vasculature. Our microrobots achieved upstream velocities of up to 1.5 μm/s and overcame blood flows of ~10 mm/s. Our results prove that microbubble-based microrobots are scalable to the complex 3D living milieu.

**Significance Statement:** Numerous brain diseases, including ischemic stroke, Alzheimer’s disease, and glioblastoma, may benefit from local and targeted therapies. Although they show great promise, microrobots have not yet demonstrated successful *in vivo* navigation inside the brain, as the challenging flow conditions and the complex 3D vascular network in the brain pose fundamental limitations. Here, we apply acoustically driven microrobots with the capacity for self-assembly and real-time navigation, including navigation against blood flow up to 10 mm/s, used for the first time inside the brain vasculature of a living mouse. The ultrasound manipulation of microrobots inside animal models provides a much-needed pathway for the advancement of preclinical research.

## Introduction

The human brain is the most complex organ that nature has created, made up of 86 billion neurons, 85 billion non-neuronal cells, and over 650 km of blood vessels (1). In spite of recent advancements in imaging technologies, brain functions remain poorly understood, and numerous brain diseases, including glioblastoma (2, 3), Alzheimer’s disease (4–6), ischemic stroke (7–9), Parkinson’s disease (10, 11), schizophrenia (12), epilepsy (13, 14), and migraine headache (15) still lack ideal therapeutic options. Brain research is a challenging field due in part to the brain complexity and its difficult access (16–20). The ability of microrobots to navigate through hard-to-reach areas can present new opportunities in medicine. These microscale automated machines can potentially navigate into the brain vasculature to assist various biological applications, such as precise drug delivery and minimally invasive surgery. In recent years, the field of biomedical microrobotics has received increasing attention; however, the *in vivo* application of these tools to brain research remains underexplored.

The brain vasculature provides a minimally invasive strategy for the administration of therapeutic compounds (3, 21–28). However, microrobots face challenging physical conditions (e.g., adverse flow, intricated vasculature network or a densely crowded heterogeneous fluidic environment) preventing them from moving towards the targeted area. Magnetic and acoustic external fields have been shown to be suitable for guiding microrobots within living tissue (29–37). Such manipulations have been seen for organs with low/no flow such as the stomach (38), urinary bladder (39, 40), liver portal vein (41), ear vasculature (42), intestine (43), lung tissue (44), cremaster muscle (45), knee cartilage (46), and cutaneous and subcutaneous vasculature (47, 48). Within the brain, only magnetic microrobots have been explored to date, and limited manipulation in situ mouse models was demonstrated with them (30). While magnetic actuation offers precise navigation, the dependence on magnetic particles limits microrobot biodegradability. Meanwhile, acoustic approaches have still not demonstrated successful microrobot trapping and navigation in physiological settings of the brain. Using acoustic manipulation, Ghanem et al. previously manipulated glass spheres that, due to their size, were restricted to large cavities such as those in the urinary bladder (40). Moreover, Joss et al. demonstrated the acoustic manipulation of single microbubbles in low blood flow zebrafish embryos (49). In the experiments of Dayton et al. inside the mouse cremaster muscle, microbubbles approached vessel walls under acoustic stimulation, but their movement was exclusively downstream (45). Finally, Lo et al. managed to acoustically trap microbubbles and navigate them inside mouse skin vessels (47).

In this article, we introduce acoustic microrobots that aim to overcome the current limitations of microscale navigation in the brain vessels. We studied the feasibility of actuating microrobots in the brain via an acoustic wave passing through a mouse skull. Our microrobots consist of aggregations of gas-filled lipid-coated microbubbles (microswarms). Under the influence of acoustic radiation, these microrobots self-assemble and simultaneously become trapped at the vessel walls, where drag forces from the blood flow are minimal. Subsequently, the microrobots move through controlled translation along the vessel wall under flow velocities of up to 10 mm/s. Additionally, we combine acoustic navigation with two-photon (2P) microscopy to achieve simultaneous real-time *in vivo* visualization of microrobots. Altogether, we present the navigation of acoustic microrobots within a living mouse brain. This method comprises an external manipulation approach that is minimally invasive, successful in complex 3D cerebral capillary networks, robust against high flow rates, and reproducible in *in vivo* environments. These results will further our understanding and the utility of ultrasound-based micromanipulation in the brains of living systems and will support novel strategies for targeted drug delivery.

## Results

### Experimental setup

The working principles of microrobot acoustic navigation in a 3D milieu were first analyzed within microfluidic vessels for later application in mouse brain vasculature. As proof of concept, we designed a microfluidic 3D vessel. Polydimethylsiloxane (PDMS) Sylgard composite was mixed with curing agent in a 10:1 ratio and poured over a copper wire with 400 μm diameter. After curing, the wire was removed, leaving behind a channel with circular cross-section. Commercially purchased microbubbles (USphere Tracer Red) were injected into the microfluidic channel and ultrasound waves were introduced via coupled piezoelectric transducers (3×3 mm dimensions and 490 kHz resonance frequency). The acoustic transducers were connected to a function generator and the whole setup was mounted on an inverted microscope and recorded with high-sensitivity cameras.

We then studied the feasibility and limitations of microrobot navigation in living mouse brains. A cranial window was implanted in the mouse to allow optical access to the brain for real-time 2P microscopy visualization (Fig. 1). Fluorescent microbubbles were subsequently injected into the mouse bloodstream via the tail vein (see *SI Appendix*, Fig. S1). The acoustic navigation set-up consisted of a piezoelectric transducer adhered to the side of the cranial window and directly coupled to the skull (Fig. 1*A*). The mouse setup was positioned on the stage of the 2P microscope used to record experimental data concerning the behavior of microbubbles inside the brain vasculature upon ultrasound activation at 35 V_PP_ and 490 kHz.

**Figure 1.**
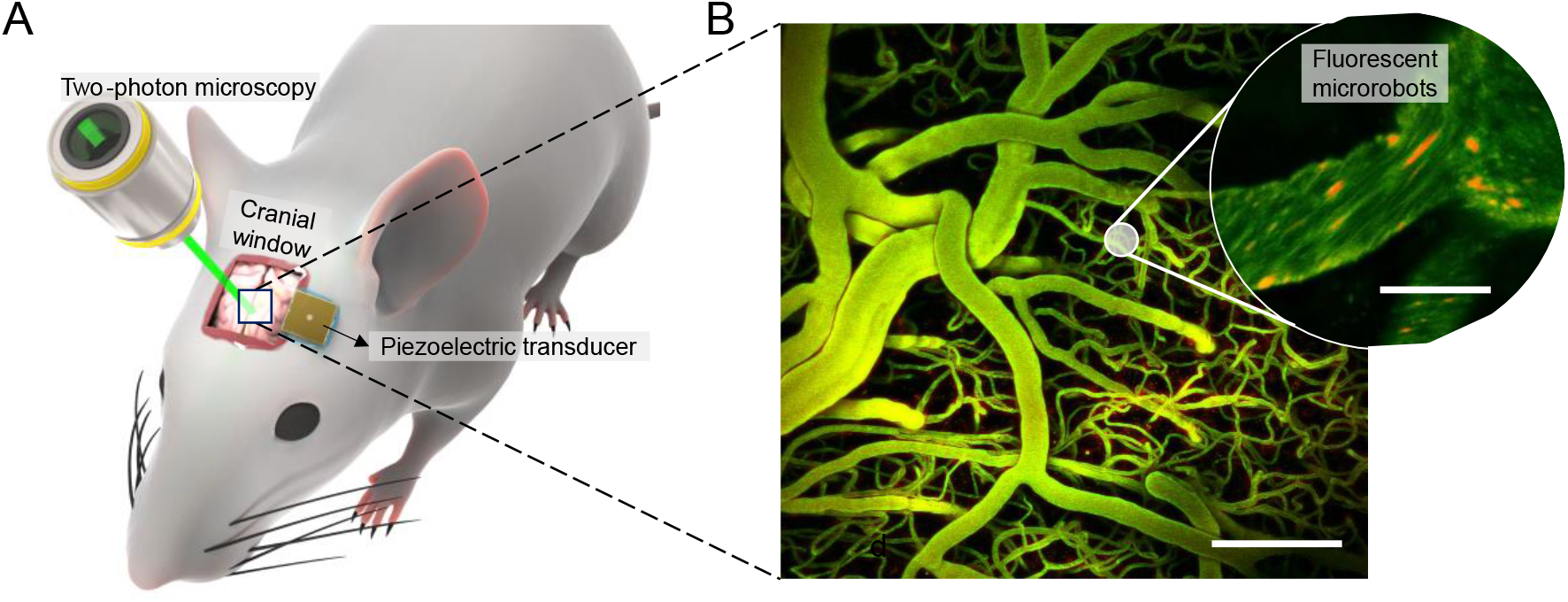
Acoustic microrobot navigation combined with real-time optical imaging. (*A*). Setup for *in vivo* studies. Optical access to the brain vasculature for 2P microscopy is provided by a cranial window. Piezoelectric transducers are attached to the skull and introduce acoustic waves directed towards the brain. (*B*). Brain vasculature as visualized by 2P microscopy (z-stack projection). Scale bar is 100 μm. In the inset, fluorescent microbubble-based swarms that will form the microrobots are seen in red. For detailed explanation of the image processing, see the Materials and Methods section. Scale bar is 10 μm.

### Transmission of sound waves from skull to brain

We investigated the external force that is exerted at the site of microrobot navigation to understand how acoustic pressure decays inside the brain. We determined the tissue depths that can be reached using acoustics and the limitations that became evident while tracking microrobots in real time with optical imaging.

We first assessed the acoustic impedance of the piezoelectric transducer and the effect of skull coupling; namely, a transducer was coupled to an *ex vivo* mouse skull and activated at 490 kHz and 35 V_PP_. We observed that upon skull connection the transducer resonance frequency suffered a subtle shift to the left and the sound wave was dampened (Fig. S2*A*). We measured the attenuation of the sound wave on its way to the brain through hydrophone 3D mapping of the acoustic pressure below the skull (*SI Appendix*). Our measurements showed pressure peaks close below each transducer, and at the z plane a decrease of the acoustic pressure along the y axis (Fig. S2*B, C*). Of note, the skull of a mouse is approximately 2 mm thick and separated from the brain by protective meningeal layers and cerebrospinal fluid (see *SI Appendix*). We found that the acoustic pressure directly below the skull is ~100 kPa while that just below the depth of the pia mater is ~85 kPa (Fig. S2*C*). Given that the depth of visualization in the *in vivo* mouse brain was 100-200 μm, the acoustic pressures at these areas range between 70-80 kPa. In our study, more than 70% of the acoustic wave was transmitted to the tissue depths in our study, where microrobots could be navigated and visualized simultaneously.

### Microfluidic validation of the acoustic 3D navigation technique

We studied microrobot formation and navigation in 3D microfluidic channels with circular cross-sections. We demonstrated that the motion-control mechanism we presented in previous studies could also move microrobots in 3D setups having arbitrary angles and relative positions of the transducer and target vessel (50). The microrobots in this study are formed by the aggregation of single gas-filled microbubbles under the influence of an acoustic wave (50). Secondary Bjerknes forces initiate the microbubble swarms, which are then translated along the vessels by primary radiation forces. It has already been established that a transducer facing the microrobots can move a bubble swarm in the direction of wave propagation. Our aim was to demonstrate that microswarm translation in mouse blood vessels is not compromised by transducer positioning: that is, microswarms will still translate when the transducer is not facing them or when the incident angle (α) of the acoustic wave is altered. For these experiments, we placed piezoelectric transducers (labeled 1-9) along a PDMS cylinder (Fig 2*A*) and activated them at 490 kHz and 15 V_PP_. We then studied the effect of various acoustic wave incident angles (α: 22.5°, 45°, 67.5°, and 90°) (Fig 2A, C). As an additional case study, we included Transducer 9, which does not face directly towards the vessel. In all cases where the transducer was facing the channel, at least 80% of microswarms formed and moved away from the transducer (Fig 2*B, E*). Transducers 2, 3, and 4, facing down at α: 67.5°, 22.5°, and 45° respectively, moved microswarms downwards along the channel length; meanwhile, transducers 6, 7, and 8, facing up at α: 67.5°, 22.5°, and 45° respectively, moved microswarms upwards along the channel length. Transducers 1, 5, and 9, placed at α: 90°, also induced microswarm formation and translation along the wall; in these cases, microswarms above and below the transducer propelled at the same rates up or down the vessel, respectively. Finally, we investigated the translation velocities of microswarms and showed that microswarm velocity decreases as α is increased up to 90° (Fig 2*D*). Taken together, our results indicate that a perpendicular channel (sound wave coming at 0°) is ideal, yielding translation velocities of up to 1.2 mm/s; however, parallel channels (sound wave coming at 90°) also display efficient microrobot translation with speeds of up to 0.4 mm/s. Thus, transducer positioning does not affect the feasibility of microswarm movement but does impact microswarm velocity.

**Figure 2.**
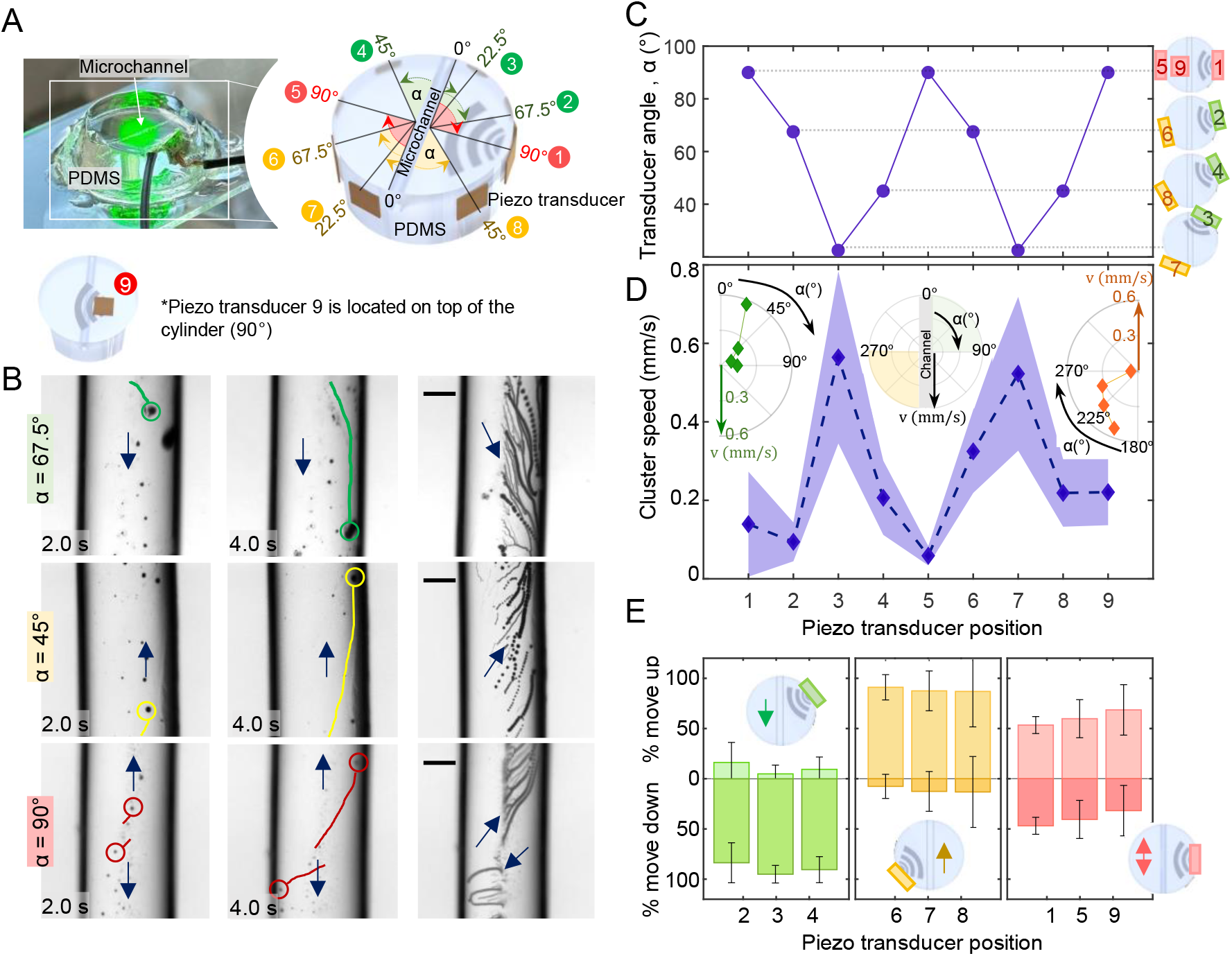
Microrobot navigation in a 3D microfluidic PDMS setup. (*A*). On the left is a photo of the experimental setup; on the right is a 3D representation of the PDMS cylinder and the positions of all transducers used for this analysis. Transducers were coupled to the side of the cylinder at various angles and labeled with numbers from 1 to 8. Transducer 9 was placed on top of the cylinder, parallel to the channel. (*B*). Microscope images of acoustic microrobot navigation under the influence of transducers 3, 8, and 9. Microrobots were manually tracked within the channel. The rightmost column is an image stack spanning four seconds of the navigation process, recorded at eight frames per second. Scales bar are 150 μm. (*C*). Plot of the angles between the channel and transducers. The relative positions of the transducers are illustrated on the right. (*D*). Plot of microswarm velocity during navigation with each respective transducer activated. Each point is the average of five measurements, connected to each other by dashed lines and the colored area represents the standard deviation. Inset subplots (in polar coordinates) represent the relation between transducer angle and swarm velocity. (*E*). Plot showing the most probable direction of swarm navigation (up or down the channel) when each transducer is activated. The bar depicts the average of seven independent measurements. Error bars are the standard deviation.

### *In vivo* self-assembly of microrobots in mouse brain vasculature

Recent studies have proposed the concept of a swarm or aggregation of units as a microrobotic approach for navigation in living systems(33, 50). However, achieving a stable and durable swarm within flowing blood has proven a major hurdle (33, 47). To test this *in vivo*, we used fluorescent microbubbles injected in the mouse blood circulation via the tail vein. These microbubbles have the capability to self-assemble into a swarm and adhere to the wall of blood vessels. We imaged fluorescent microbubbles in the cortex of living C57BL/6 mice by using intravital 2P microscopy. After allowing the microbubbles to circulate freely for 5-10 minutes, an acoustic transducer coupled to the mouse skull was activated at 490 kHz and 35 V_PP_. The acoustic signal was continuously monitored by an oscilloscope.

In response to acoustic activation, we observed that microbubble clusters started forming at vessel walls inside the brain (Fig. 3*A,B*, see *SI* Movie S1). At these boundary regions, microbubbles encountered low drag force from the blood flow, which allowed acoustic radiation forces to dominate, resulting in microbubble aggregations. Importantly, microrobots were able to endure physiological flow forces as long as the acoustic signal was activated. Once the transducer was deactivated, the microswarms disassembled and the loose bubbles followed the natural downstream blood flow. We characterized the evolution of swarm size inside blood vessels over time and we were able to distinguish swarm sizes ranging from 8 to 40 μm on the basis of the time of acoustic activation. We observed that as hemodynamic forces brought more microbubbles to the activated site, the clusters continued to grow, resulting in swarms of size up to 40 μm. Large swarms tend to grow fast within the first seconds until they reach a saturation state, while smaller swarms tend to keep growing over time (Fig. 3*C*). The variability in microswarm formation speed comes in part from the complexity of the brain vasculature network and the resulting relative position of each blood vessel to the transducer. We validated these data in our microfluidic devices by showing that microbubble velocities are affected by relative position and distance to the acoustic transducer.

**Figure 3.**
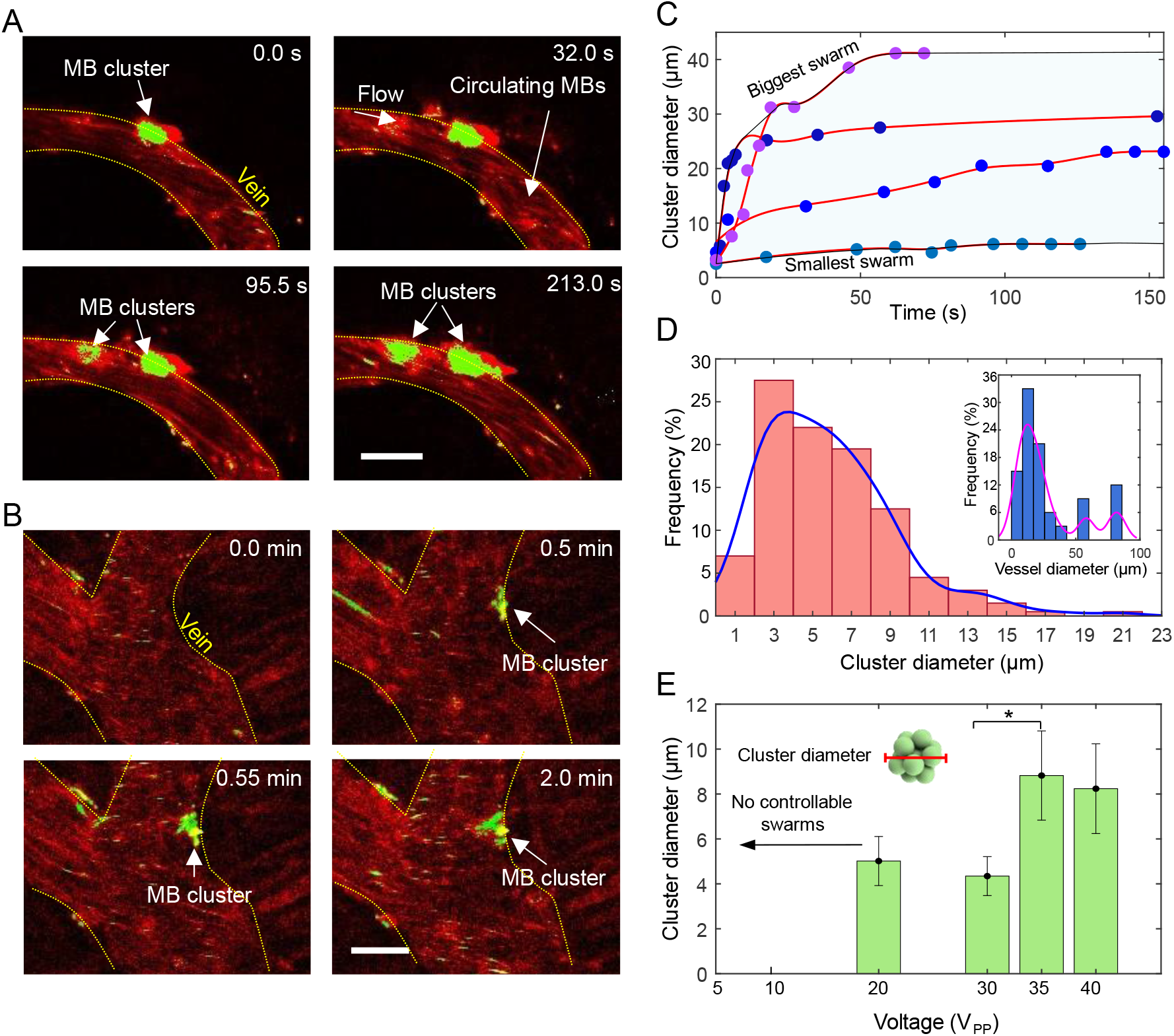
Microbubble-based swarm formation in mouse brain vasculature. (*A, B*). Microbubbles aggregate over time and adhere to the walls of blood vessels under an acoustic signal of 490 kHz and 35 V_PP_. After image processing (see Materials and Methods), the microswarms have been colored in green while non-responding single microbubbles that flow downstream have been colored in red. Scales bar are 50 μm. (*C*). Evolution of swarm formation and growth under a signal of 490 kHz and 44 V_PP_. Plot shows cluster diameter over time for four different microswarms at four different locations around the brain vasculature. Each point is one microswarm at one time point. Swarm size increases over time until reaching saturation. We colored in light blue the area of possible swarm size evolution, as observed during experiments. (*D*). Plot showing the cluster diameter distribution observed at an acoustic excitation of 490 kHz and 35 V_PP_. A total sample of 200 clusters was analyzed. The inset at top right illustrates the vessel size distribution observed during experiments. A total sample of 39 vessels has been studied. This represents the vessel sizes wherein we could achieve microrobot navigation. Fitting lines have been drawn to summarize the tendency for each plot. (*E*). Plot showing cluster diameter versus voltage (at a constant acoustic frequency of 490 kHz). Higher voltage results in bigger microbubble clusters, on average. Each average was determined from 25 measurements and the error bars represent the standard deviation.

In addition, we observed that more frequent swarm formation occurred at lower flow velocities suggesting that microbubbles aggregation is influenced by blood flow rates. During the experiments, we mostly observed increased microbubble formation in vessels with low flow (veins and capillaries with diameters of 10-40 μm) compared to vessels with high blood flow velocities (arteries and arterioles Fig. 3*D*). Meanwhile, the microbubble clusters themselves were mostly between 2 and 10 μm in diameter, with larger swarms found in larger vessels (Fig. 3*D*). Although these analyses were performed under constant acoustic conditions of 35 V_PP_ and 490 kHz, we also studied the effect of decreasing the applied voltage from 40 to 20 V_PP_; this halving of voltage resulted in the average cluster size also being reduced by half (Fig. 3*E*).

Collectively, these data provide the first characterization of the dynamics of microswarm formation in a brain blood vessel and constitutes a first step towards the implementation of microrobot navigation *in vivo*.

### Navigation of microrobots inside the mouse brain vasculature

The presence of flow inside blood vessels is not only a challenge for microrobots formation but also for *in vivo* navigation. The propulsion of microrobots is normally hindered when in flow, and navigation against flow represents an even bigger hurdle. In our experiments, we want to prove that the navigation capabilities of our acoustic microrobots are maintained when moving to an *in vivo* flow environment.

When microbubbles aggregate and form swarms, there is an increase in their overall volume, which in turn increases the acoustic radiation forces that are exerted on them. We observed that the process of navigating our microrobots is marked by two behaviors: (1) microswarms continue attracting each other and grow even during navigation (Fig. 4*A*) and (2) the swarms are propelled along the vasculature (Fig. 4*B,C*, see *SI* Movie S2, S3). The challenge of this task is significantly higher when the microswarms need to be moved in the direction opposing blood flow, thus in this work we focused on the upstream movement of microswarms. Under acoustic activation at 490 kHz and 35 V_PP_, we achieved upstream movement of microbubble aggregations at velocities of up to 1.5 μm/s. Compared to our *in vitro* results, the speed of microrobots within a cerebral vessel was lower than the speed of microrobots in the microfluidic channel. We suggest that these differences are due to the acoustic attenuation by the meningeal layers, brain parenchyma and blood composition. Further, we confirmed that microbubble formation and navigation was mostly successful inside veins and capillaries with vessel diameters between 10 to 40 μm, and the microswarms were 3 to 10 μm in size. Importantly, the microswarms overcame blood flow velocities of up to 10 mm/s (Fig. 4*D,E*).

**Figure 4.**
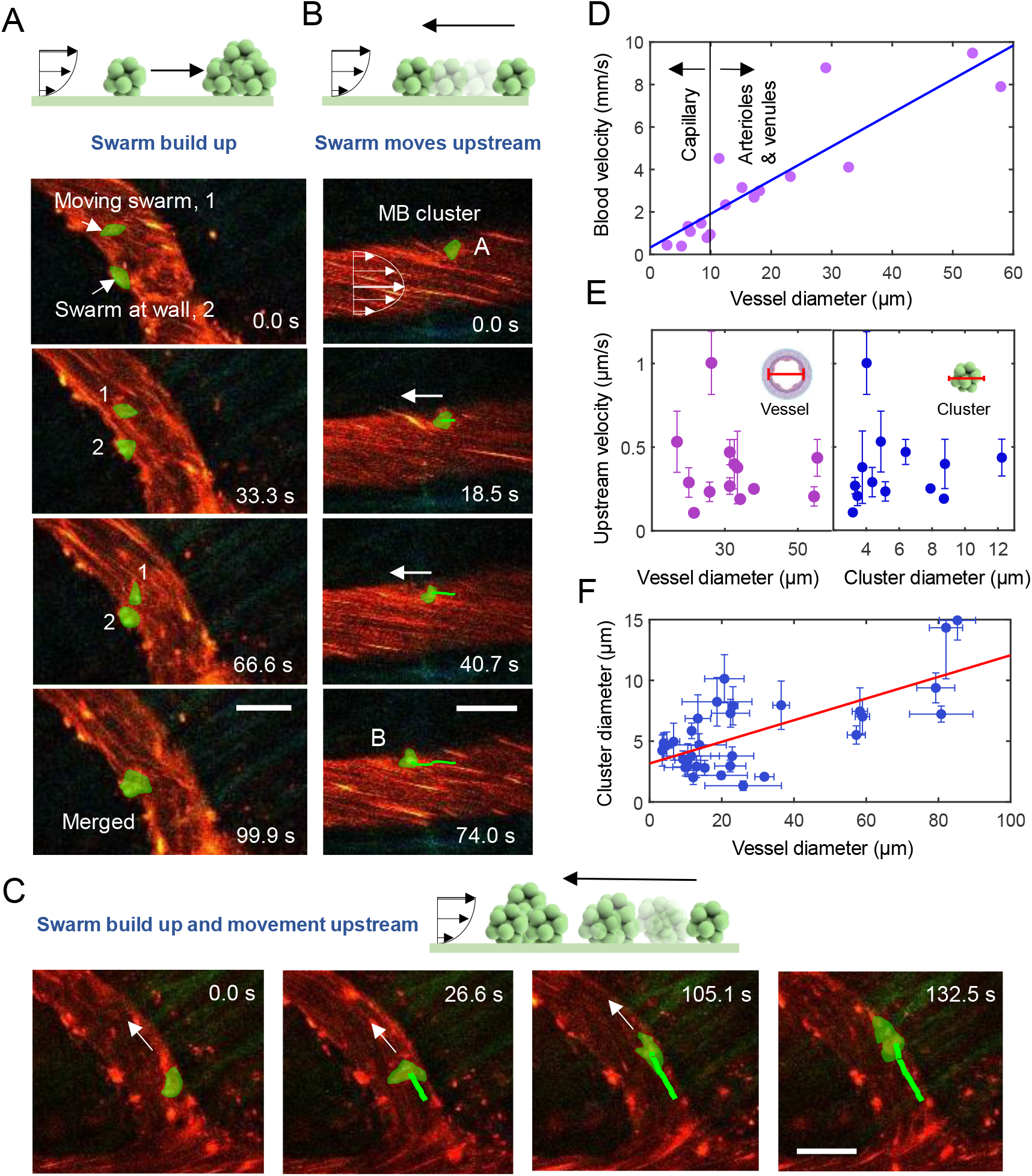
Microswarm upstream navigation in mouse brain vasculature. (*A*). At the top, the schematic represents the first possible situation of microswarm navigation downstream and simultaneous growth. In the images below, we observe a microswarm (colored green by manual overlay) navigating in a controlled downstream trajectory (not washed away by the background blood flow) and finally being attracted to a second microswarm that was already at the wall. The acoustic signal was 490 kHz and 35 V_PP_. Scale bar is 30 μm. (*B*). At the top, the schematic represents the second possible situation of microswarm navigation upstream without growth. In the images below, the movement of a microswarm (green overlay) is manually tracked along a vessel wall and against blood flow direction. The acoustic signal was 490 kHz and 44 V_PP_. Scale bar is 30 μm. (*C*). At the top, the schematic represents the last possible situation, microswarm navigation upstream and simultaneous growth. In the images below, the movement of a microswarm (green overlay) is manually tracked against the flow, aggregating more bubbles along the way (see Movie S3). The acoustic signal was 490 kHz and 35 V_PP_. Scale bar is 30 μm. (*D*). Plot of the blood flow velocities regularly found in the mouse vasculature, for those vessel diameters for which we observed microswarm formation and navigation. We have drawn a linear fit line with R^2^=0.7839. (*E*). Plot indicating the observed upstream velocity of microswarms versus the vessel diameter and swarm diameter. The acoustic signal was 490 kHz and 35 V_PP_. Each point is the average of five measurements and error bars are the standard deviation. (*F*). Correlation between vessel size and cluster diameter, showing the tendency to bigger swarms in bigger vessels. Each point is the average of five measurements and error bars are the standard deviation. R^2^ for the fitting line is 0.4719. The acoustic signal was 490 kHz and 35 V_PP_.

Our results suggest that microswarm propulsion in the brain vasculature is directly affected by two phenomena. First, microrobot speeds are compromised by internal flow conditions. Second, the acoustic radiation force on the swarm scales proportionally with its volume, with the higher force experienced by bigger swarms translating to faster navigation. Altogether, these data reveal how large swarms form in larger vessels (Fig. 4*F*) and why microswarms displayed similar upstream velocities independent of swarm size or vessel size (Fig 4*E*). Collectively, our findings prove the successful acoustic navigation of microrobots upstream against *in vivo* physiological flows.

## Discussion

In this study, we have validated the feasibility of acoustic microrobots to navigate within the *in vivo* mice brain vasculature. First, the results of these experiments demonstrate that microbubbles working in an acoustic wave exhibit the same behavior *in vivo* as those in microfluidic channels. Second, we revealed that microrobots navigation is scalable from single vessels to complex vascular networks such as those in the brain. Through these studies, we have shown that acoustic damping from the skull does not compromise microrobot navigation within the pial vasculature of the brain. We used 2P microscopy for the real time imaging of microrobots, which enabled the study to be performed at superficial tissue layers. Additionally, we measured acoustic pressures below an ex vivo skull and showed that microrobot control at deeper regions is also attainable.

Notably, through our experiments in microfluidic device, we showed that the position of an acoustic transducer determines the magnitude of microbubble response. This implies that the transducer positioning impacts how fast the swarms self-assemble and move within a channel. Nevertheless, in all cases microbubbles move along the vessel wall and in the direction of wave propagation, under the constrains of the boundaries of the vessel. In this study, we translated the results from a microfluidic context to *in vivo* conditions. Note that the experiments that we performed in mice were done in every case at a similar tissue depth but at different locations around the head of the mouse, resulting not only in different distances from the transducer, as well as different orientations with respect to it, introducing variability to the acoustic microrobot control (see *SI Appendix*, Fig. S3). This variability is represented in Fig. 4C, where microrobots are displayed at a range of upstream velocities. Furthermore, we performed a comparison between the vessel sizes where we manipulated microrobots and their average blood flow velocities, showing successful microrobot formation and navigation in flow velocities that go up to 10 mm/s. These results reassert the capabilities of acoustic microswarms to work under *in vivo* physiological conditions.

While we believe our study provides a significant step toward the use of microrobots in brain research, a few technical issues must also be taken into account. Ultrasound is a non-ionizing wave that penetrates the body with minimal invasion, but it can still trigger unwanted effects, such as heating. We performed additional measurements of temperature over time at a fixed spot below the skull (*SI Appendix*, Fig. S5), and we observed a temperature increase that should be considered in further studies and biocompatibility analyses.

Thus, we demonstrated controlled microrobot self-assembly and navigation inside living brain vasculature. The microswarms we developed show promising potential for use as drug carriers for drug targeting and delivery applications.

## Materials and Methods

### In vitro microfluidic set-up

We fabricated *in vitro* channels mimicking the size and mechanical properties of blood vessels via PDMS Sylgard composite. For its fabrication PDMS Sylgard base was mixed with curing agent in a 10:1 ratio, followed by degassing and pouring into a mold with a wire inside. The wire used was 400 μm in diameter and was previously coated with silane. The PDMS was cured during 1 hour at 85°C. After curing, the wire was pulled out of the PDMS. Piezoelectric transducers were coupled to the sides of the PDMS block. A square signal was applied to the piezo elements with a function generator (Tektronix AFG3000) with an amplitude varying from 0.1 V_PP_ to 15 V_PP_. The imaging was realized with an inverted microscope (Zeiss 200m), and images were recorded with a high-quality camera (Zeiss AxioCAM 305).

### Microbubbles

Microbubbles used for *in vitro* and *in vivo* experiments were commercially purchased from USpheres. We used USpheres Tracer-red; these bubbles have a size distribution of 1.1 – 1.4 μm and a concentration of ~2.5×10^10^ bubbles/ml (see *SI Appendix*, Fig. S1).

### Animal preparation

All animal experiments were approved by the local veterinary authorities in Zurich and conformed to the guidelines of the Swiss Animal Protection Law, Veterinary Office, Canton of Zurich. For *in vivo* experiments C57BL/6 mice aged 8-12 weeks were used. Cranial window implantation was performed as described earlier(51). For the experimental procedure, mice were anesthetized with isoflurane (induction 4% and maintenance at 1.2%, supplied with 300 ml/min 100% oxygen) and were kept at 37°C with a homeothermic blanket (Harvard Apparatus). Fluorescent microbubbles and FITC-Dextran (70 kDa, Sigma-Aldrich, cat. No. FD70S) were injected into the blood stream of the mouse via tail vein injection. The piezoelectric element was placed with superglue to the skull, next to the cranial window (Fig. 1A). Transducers were connected to an oscilloscope for data verification and to a function generator Tektronix. This set-up was mounted onto the stage of the 2P microscope, for real-time tracking of microrobots. More details on pressure measurements, transducer characterization and temperature recording can be found in *SI Appendix* and Fig. S2, S4 and S5.

### Image processing

We implemented image processing to the experimental results recorded via 2P microscope during *in vivo* experiments. Microswarms (bubble aggregations) exhibited high fluorescent intensities; thus, we utilized an intensity-based thresholding on each image, to create a mask that only identifies microswarms, excluding single microbubbles within blood flow (see Fig. 3). For the results on Fig. 4, this mask was defined manually, due to poor resolution while the swarm is moving. The swarm mask was set as green and overlaid on the original image, where single microbubbles were tinted in red.

### Statistical analysis

All the statistical analysis has been executed using MATLAB and MATLAB curve fitting toolbox. All the results presented in the figures depict the average of at least 5 independent measurements and its associated standard deviation. We have used R^2^ to assess curve fittings. A total sample of 200 microbubble clusters and 39 blood vessels was studied during the experiments. More details on statistical analysis are found in the corresponding figure captions.

## Acknowledgments

This project has received funding from the European Research Council (ERC) under the European Union’s Horizon 2020 research and innovation programme grant agreement No 853309 (SONOBOTS) and ETH Research Grant ETH-08 20–1. The authors thank Dr. Mehmet Fatih Yanik and Dr. Mehmet Ozdas for helpful contribution on acoustic pressure measurements.

