## Supplementary information for "Acoustic trapping and navigation of microrobots in the mouse brain vasculature"

**This PDF file includes:**

Supporting text
Figures S1 to S5
Legends for Movies S1 to S3
SI References

**Other supporting materials for this manuscript include the following:**

Movies S1 to S3

**Supporting Information Text**

**Extended Materials and Methods**

**Microbubble composition and fluorescence.** During all the experiments we have used commercially available gas filled microbubbles (USpheres). These microbubbles are formed with a phospholipid outer layer that contains an inert gas on the inside, specifically, perfluoro propane (C3F8), see Fig. S1.

The microbubbles are received inside a glass vial, with a saline solution. We used the UltraMix Agitator (Bio Sonics Trust) for 40 seconds for microbubble production. The final result is a vial with 0.8 ml and a concentration of ~2.5x1010 bubbles/ml, as detailed by the provider. The size distribution of these bubbles is 1.1 – 1.4µm.

For in-vivo experiments, we used 2photon as real time imaging technique with high resolution. In order to visualize the microbubbles with 2-photon microscope, we used USpheres combined with a fluorophore, which are called USpheres Tracer- Red. The fluorophore has been incorporated inside the phospholipid layer, as shown in Fig, S1. Tracer FD-Red uses a dye with excitation at 549nm and emission at 565nm.

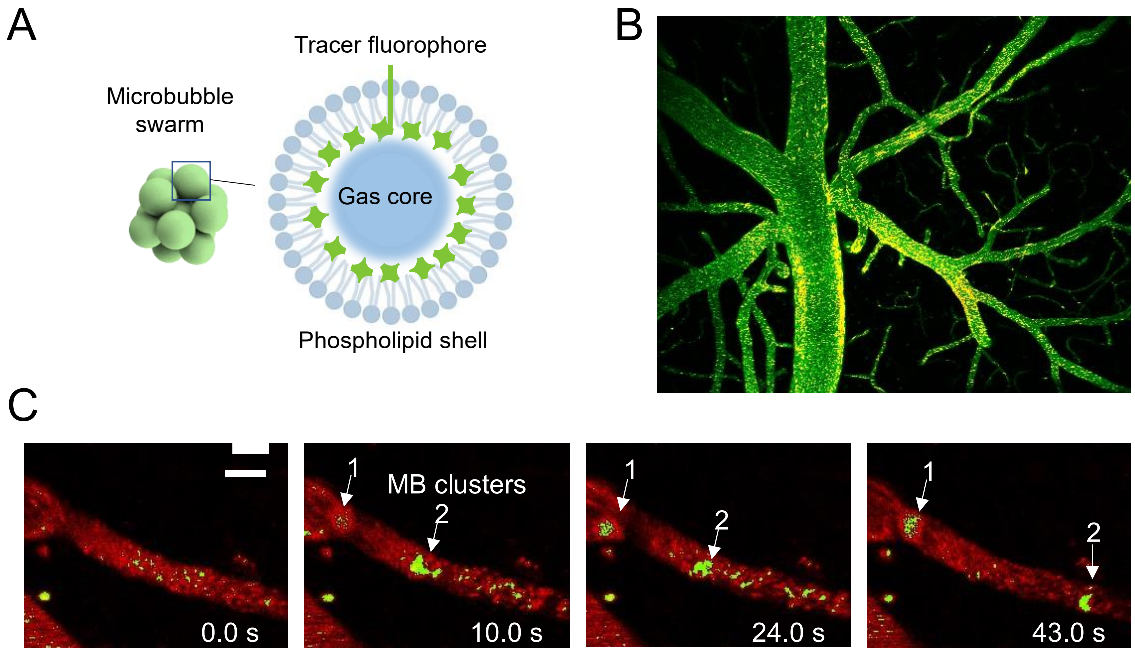


**Fig. S1.** Microrobot and microbubble composition inside the pial vasculature of the brain. *(A)* Microrobots are composed by an aggregation of microbubbles. We used USpheres, which are microbubbles with a lipid coating and perfluoro butane gas in the inside. We manipulated and visualized microbubble swarms inside the pial vasculature of the brain. *(B)* 2 photon microscope image of the pial vasculature of mouse brain. Bright green dots are the microbubbles flowing inside the vasculature. *(C)* Images from the 2 photon microscope visualization of the pial vasculature of the brain when ultrasounds are activated. Microswarms have formed and they move along the vessel walls. The microswarms have been colored in green; see Image processing in Materials and methods section of the manuscript.

**Measurement of the intracranial pressure field and piezo transducer characterization.** The characterization of the acoustic field at the excitation frequency 490 kHz has been done in a deionized and degassed water-filled water tank with a 2.5 mm needle hydrophone (HNR, HONDA) coupled to a 20 dB pre-amplifier (AH-1100 Preamplifier, HONDA). The temperature of the water bath was set to 25 °C. The control computer was driving 10 cycles, 67 mV pulse sending from the function generator at a PRF of 100 Hz provided by a digital oscilloscope (PicoScope, 5044D). At the same time, the hydrophones signal output was recorded through the digital oscilloscope for every spatial point in the pre-defined grid.

All the piezo transducers used in this document have been commercially purchased through STEMINC and referenced as SMPL3W3T05410. Their dimensions are 3x3x0.55 mm. An impedance analyzer has been used to measure the impedance of the transducer given an excitation frequency. The measurements consisted of a frequency sweep from 100 kHz to 1 MHz with a 1 kHz step and a 3 ms delay between each measurement. We use 490 kHz as resonance frequency for our experiments.

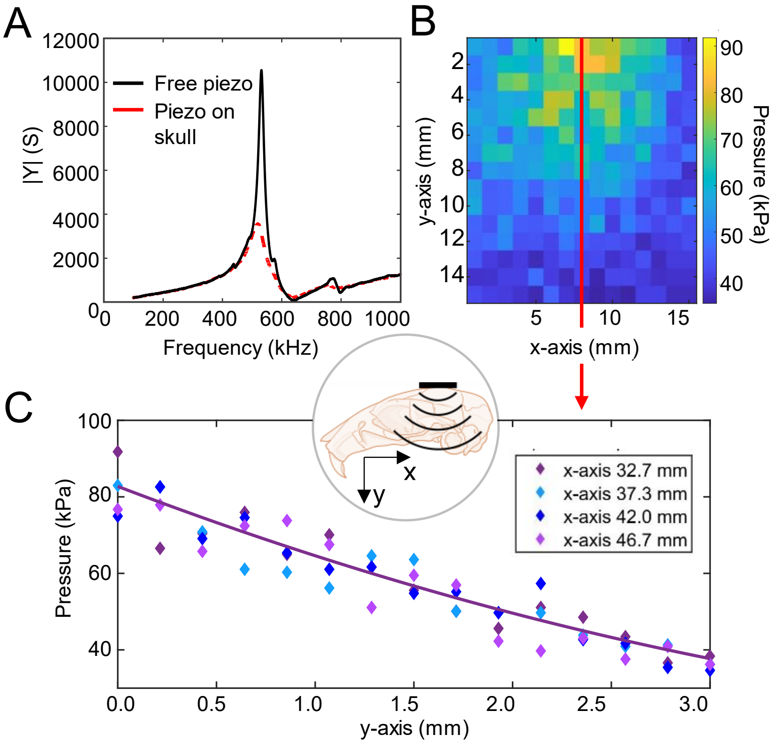


**Fig. S2.** Acoustic pressure measurements within an *ex vivo* mouse skull. (*A*). Impedance analysis of the piezoelectric transducer (490 kHz resonance frequency, as detailed by the provider) with and without skull coupling. (*B*). 2D map of the acoustic pressure below the skull upon transducer activation at 490 kHz and 45 $V_{PP}$. For more details on the measurement procedure, see Materials and Methods. (*C*). 1D acoustic pressure along the y axis of the skull at different x values. A general attenuation decay is observed, reflected by the best-fit curve. $R^2$ =0.8802 was used for fitting assessment. Inset shows a schematic of the mouse skull, definition of x and y axes, and transducer position during measurements.

The skull of mice has between 1-2 mm depth and it’s followed by protective meningeal layers and cerebrospinal fluid before reaching the brain tissue (5, 6). The main role of meningeal layers is to protect our brain from physical trauma, thus they act as shock absorbers(6). Although these layers are very thin, they will still contribute to the attenuation of the acoustic wave. We need to consider that the acoustic pressures we measured below the skull, do not include the contribution from the meningeal layers. Even so, we studied the feasibility for the remaining acoustic pressure to activate microrobots.

**Positioning of piezoelectric transducer on top of a mouse skull.** The chosen living systems for experiments were mice, thus the acoustic set-up needed to be adapted to the small size of these animals. We used piezoelectric transducers of 3x3 mm dimension and 490 kHz resonant frequency and we investigated the attenuation of the sound wave on its way to the brain. The skull was cleansed such that no soft tissue was adhered to it, and we coupled a piezoelectric transducer to the top side of the skull. We activated the transducer at the excitation frequency of 490 kHz. Although in the present, there is a broad use of focused ultrasound techniques that can place the transducer separated from the skull (1–4); here we are investigating the acoustic response of microrobots in the absence of a focusing force; thus the transducers had to be coupled directly to the bone.

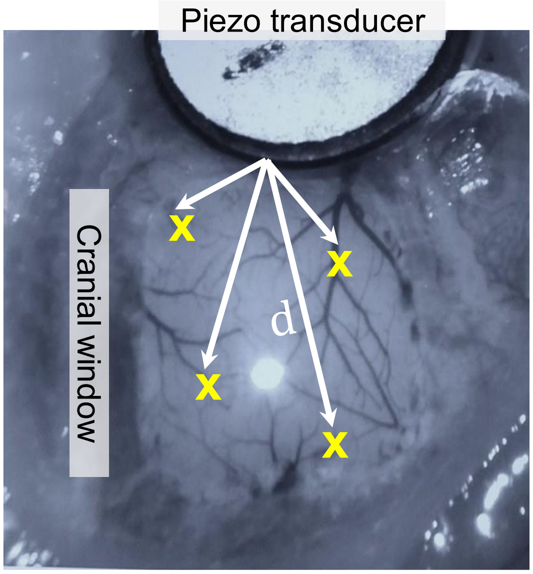


**Fig. S3.** Variability introduced from distance and orientation between transducer and blood vessel. Experiments in vivo were performed every time at similar tissue depths, however, we screened the whole cranial window. In this image we see the cranial window from above and a piezoelectric transducer coupled on the top side. Thus, in each experiment we have a different distance ‘d’ (distance to the transducer) and different angle ‘α’ ( angle between acoustic wave propagation and blood vessel orientation). This variability is translated into different microswarm formation rates and different swarm speeds inside the blood vessels. The experiments we performed didn’t track the position of the field of view; thus we work with this intrinsic variability.

**Simulations of acoustic pressure below a mouse skull.** A numerical model of a rodent skull and brain was designed to evaluate the spatial distribution of the acoustic field inside the cortex of the animal. These data would provide an overview for the upcoming Vivo study. The numerical study also allows testing of different parameters and especially different range of powers. By computing mechanical and acoustic power dissipation, it is also possible to estimate the temperature increase inside the brain.
To have a better overview of the acoustic field map below a mouse skull, it has been decided to pursue the study with a 3D model, see Fig S4. The skull geometry is taken from a 3D scan of a juvenile female mouse made available through ONE Core facility39. We added to the geometry 3 5*5*0.4 mm piezo elements and a fluid-filled cavity representing the brain (COMSOL 6.0). An

appropriate mesh for the finite element analysis was built based on the physics used in the finite element analysis software.

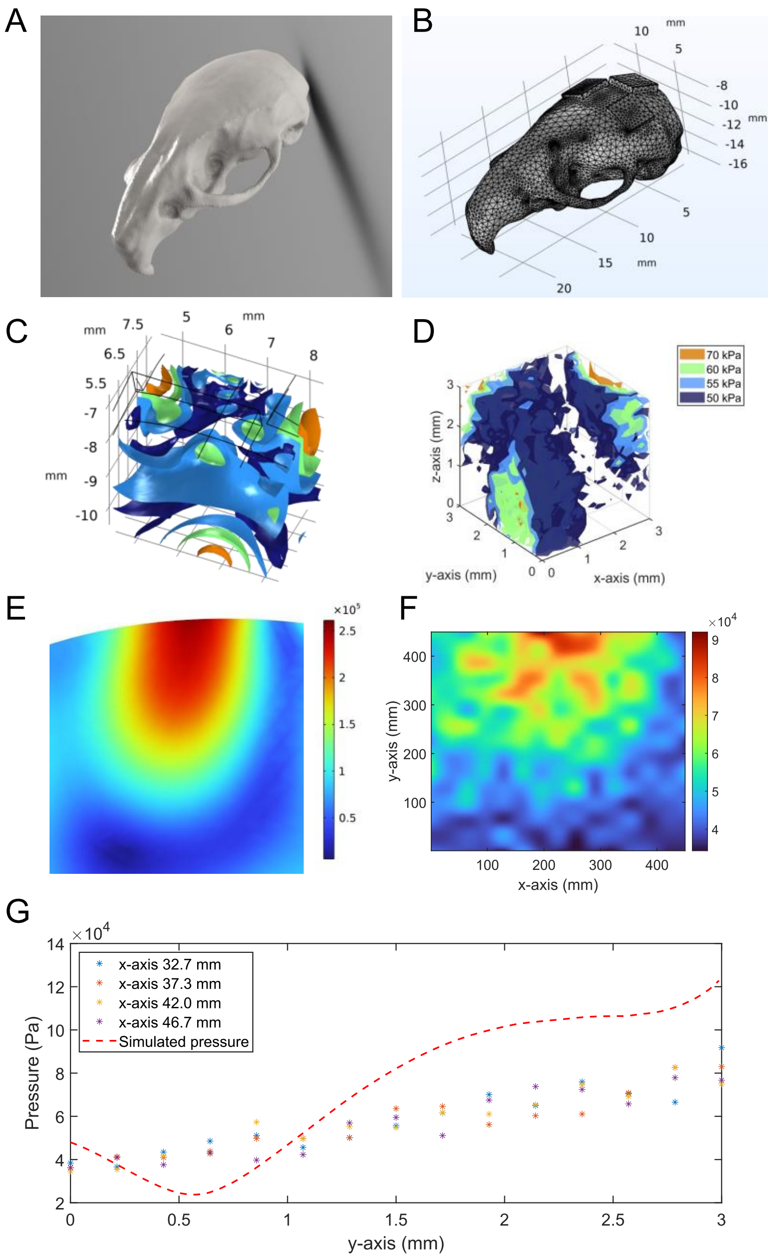


**Fig. S4.** Acoustic pressure simulation results and its comparison with experimental measurements *(A)* Skull scan .stl file from ONE Core. *(B)* Skull import in COMSOL, with three piezo elements added on top of the skull, and meshing of the whole geometry for finite element analysis. *(C)* Intracranial pressure field computed by the numerical simulation below three piezo transducers. *(D)* Interpolated intracranial pressure field experimentally measured below three transducers. *(E)* X-Y plane of the simulated pressure field at an antinode. *(F)* X-Z plane of the measured pressure field at an antinode. *(G)* Plot comparing the evolution of the simulated versus measured pressure field along the y-axis and centered at the maximum pressure along the x-axis.

The numerical model uses 3 physics modules (acoustic, mechanic, and electrostatic) for 3 different domains (skull, brain, and piezo elements). Piezo elements were modelized as PZT-5A material and simulated by coupling mechanic and electrostatic physics module. They were numerically excited at a frequency of 490 kHz and an amplitude of 25 $V_{PP}$. The propagation of the acoustic waves due to piezo transducer motion has been computed with the mechanic module. The displacements generated are below the micrometer range and the strains are lower than $10^{-3}$ , thus a linear elastic material model complemented with damping has been chosen to model the skull bone. Lastly, the acoustic field in the skull cavity has been computed with the acoustic module and appropriate damping. The skin has not been taken into account here as in our Vivo experiment the mouse cranial epidermis was removed to build a cranial window.

Our numerical model was compared with a 3D scan (3*3*3 mm) of the intracranial pressure field in an Ex Vivo rodent, see Fig. S4. As in the simulation, the three piezo transducers were excited with 10 cycles pulse at a frequency of 490 kHz and an amplitude of 25 $V_{PP}$. A particular pattern of 3 pressure antinodes has been observed analogous to the results output by the simulation. The order of magnitude are quite similar. Lastly, if we compare the measured and simulated pressure distribution along a y-axis line, we observe that absolute pressure amplitude ranges from 70 kPa to 100 kPa at the interface between the skull and brain cavity, Fig. S4.

**Measurement of temperature below ex vivo mouse skull.**

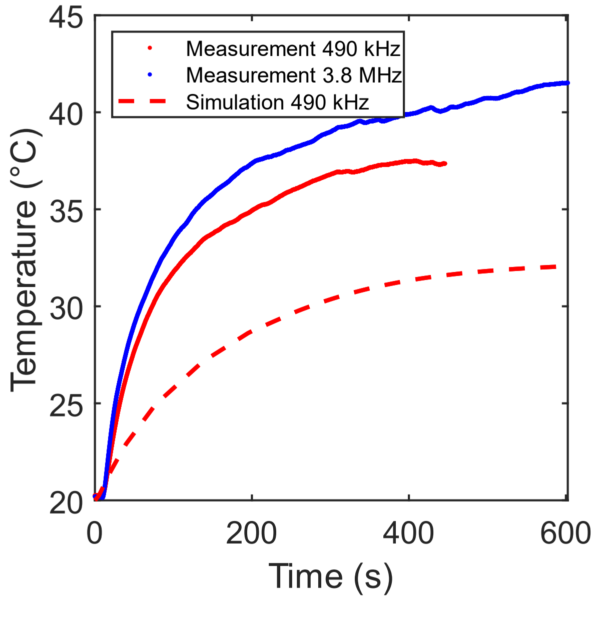


**Fig. S5.** Plot showing the increase of temperature caused inside water due to acoustic activation on top of an ex vivo skull. We show measurements next to simulation results.

The recording of the temperature in an ex vivo mice head has been done while three piezo transducers were glued at the same position as in the in vivo experimental setup. A type K thermocouple probe was gently inserted into the brain by the ventral side of the skull. A thermocouple data logger (TC-08, Pico technologies) was used to log and record the thermocouple output on a computer. The three piezo elements were excited with a square signal at the frequency of 490 kHz and with 20 $V_{PP}$ amplitude until the temperature reached equilibrium.

**Movie legends**

**Movie S1 (separate file).** Microswarm navigation to the wall and growth

(27.9 MB)

Within this movie we have two separated videos, they correspond to the results shown in Fig. 3 *A,B*. In both videos we can see microbubbles aggregating into bigger swarms at the vessel wall of a mouse brain. In both vessels there is a prominent blood downstream flow; nevertheless microbubbles are unaffected by it, and they aggregate and stay at the wall. We applied acoustic excitation at 490 kHz and 35 V peak-to-peak. The video was captured at 0.74 frames per second (fps) and played at 10 fps.

**Movie S2 (separate file).** Microswarm upstream navigation inside in-vivo brain vasculature

(16.2 MB)

The video corresponds to Fig. 4*B*. The video shows the upstream navigation of a microswarm along the vessel wall of a mouse brain. We applied acoustic excitation at 490 kHz and 44 V peak-to-peak. The video was captured at 1.5 fps and played at 10 fps.

**Movie S3 (separate file).** Microswarm upstream navigation and simultaneous growth

(16.0 MB)

The video corresponds to Fig. 4C. The video shows the navigation of a microswarm upstream along a vessel wall. On the way, the microswarm attracts more microbubbles and grows. The final microswarm is not only upstream, but also bigger than before. We applied acoustic excitation at 490 kHz and 35 V peak-to-peak. The video was captured at 0.74 fps and played at 10fps.
